# Deciphering Cannabidiol Neuroregulatory Role in Addiction Pathways: A Systems-Level Comparison with THC via Intrinsic Network Pharmacology

**DOI:** 10.1101/2025.07.26.666970

**Authors:** Hemanth Kumar Manikyam, Sunil K. Joshi

**Author notes:** **Correspondence:** Hemanth Kumar Manikyam, Email: [ ]; Address: Faculty of Science, Department of Pharmacology, North East Frontier Technical University, Aalo-791001, Arunachal Pradesh, India: Professor Chalapathi institute of Pharmaceutical Sciences, Acharya Nagarjuna University -Guntur: India.

## Abstract

Cannabidiol (CBD), a nonpsychoactive phytocannabinoid from Cannabis sativa, has demonstrated potent neuroprotection in a number of neuropsychiatric disorders. Unlike Δ9-tetrahydrocannabinol (THC), which acts to stimulate the brain’s reward circuitry through CB1 receptor activation, CBD seems to stabilize several neuroregulatory circuits without causing addictive reinforcement. Yet a mechanistic systems-level distinction between these two cannabinoids is relatively unexplored. This research uses an eight-layer Intrinsic Network Pharmacology (INP) system to model and compare the molecular effects of CBD and THC on addiction-related networks, combining computational modeling and network pharmacology. The INP system includes molecular trigger mapping, feedback loop dynamics, redox balance, immune signaling, autophagy repair, therapeutic fit, dynamic simulations, and multi-target synergy overlays. In this model, we replicated the action of CBD on major molecules such as Nrf2, ROS, IL-6, dopamine (DA), D2 receptor, and BDNF. Ordinary differential equation (ODE) models were employed to model time-dependent transitions, whereas Boolean logic models simulated binary molecular switches. Comparative reward pathway diagrams were drawn to represent divergent neuropharmacological cascades triggered by CBD and THC. Our simulations show that CBD triggers redox restoration, inhibits inflammatory cytokines, and induces dopaminergic and synaptic stability. Withdrawal simulation also showed that relapse of the partial system could be triggered if CBD is withdrawn suddenly, substantiating its status as a homeostatic stabilizer and not a suppressive one. Network pharmacology fit scoring exhibited CBD’s high concordance with antioxidant and neuroplasticity targets, whereas THC mostly mapped onto reward-amplifying nodes. Though these results are hypothesis-driven and computational, they also offer a useful framework for the experimental validation. This work illustrates the utility of INP-based simulations in deciphering polypharmacological drugs such as CBD and presents a scalable paradigm for assaying future candidates for neurotherapeutic use in the study of addiction.

## 1. Introduction

Cannabis sativa has been a plant of medicinal and sociopolitical importance for centuries, containing more than 100 phytocannabinoids that play complex physiology. Of these, Δ9-tetrahydrocannabinol (THC) and Cannabidiol (CBD) have been most researched because of their unique neuropharmacological profiles [1]. THC is known to be the major psychoactive compound that produces cannabis’s euphoric and reinforcing effects. It engages effectively with CB1 receptors in the brain’s mesolimbic dopamine system, stimulating reward circuits and possibly playing a role in the onset of cannabis use disorder (CUD) [1]. CBD, on the other hand, has no psychoactive effects and has a complex pharmacology that involves allosteric modulation of cannabinoid receptors, serotonergic activity, antioxidant action, and even inhibitory action on drug-seeking [2].

As global legalization and medicinal use of cannabis and CBD-containing products continue to grow, it is critical to be able to differentiate between those compounds that enhance addiction susceptibility and those with therapeutic potential for anti-addiction [1,2,3]. The role of THC in the reinforcement of dopaminergic circuits has been well documented. It stimulates G-protein-linked CB1 receptors to enhance dopaminergic activity in the ventral tegmental area (VTA) and the nucleus accumbens (NAc), key elements of the reward pathway [3,4]. This activation model’s classic addictive drugs such as opioids and cocaine, which also increase dopamine release and strengthen acquired reward behavior [4].

CBD, by contrast, is structurally homologous with THC but pharmacologically different. It has very low binding affinity to CB1 and CB2 receptors [5,6] but rather serves as an inverse agonist at CB1, potentially lowering the psychoactivity of THC in combination. In addition, CBD acts on a wide variety of molecular targets such as TRPV1 (transient receptor potential vanilloid 1), 5-HT1A (serotonin receptor subtype), GPR55, PPARγ, and adenosine receptors [5]. This wide target profile supports CBD’s neuroprotection, anxiolysis, anti-inflammatory, and immunomodulatory actions—making it a molecule of significant interest in addiction neuroscience, epilepsy, and psychiatric illness [5].

Current preclinical and early clinical trials have demonstrated that CBD can decrease drug-seeking behavior, decrease craving, and modulate dopaminergic transmission [6]. It seems to restore homeostasis in substance-abuse-disrupted neural circuits, such as normalizing glutamate and dopamine neurotransmission, inhibiting pro-inflammatory cytokines including TNF-α and IL-6, and triggering antioxidant pathways, particularly Nrf2 signaling. Further, CBD has been found to increase BDNF (brain-derived neurotrophic factor) expression, which underlies synaptic plasticity and neuronal survival—both of which are characteristically deficient within the addicted brain.

Notwithstanding these nascent findings, there still exists a dearth of systems-level models integrating the long-term physiological effect of THC and CBD upon addiction-relevant networks [7]. Most researches study isolated receptors or behaviors, not considering the multilayered, dynamically interacting biological systems underlying addiction. Bridging this deficiency, we present a systems pharmacology and computational strategy called Intrinsic Network Pharmacology (INP) that simulates molecular and regulation cascades through eight layers of biology: trigger/collapse nodes, feedback loops, redox dynamics, immune signals, autophagy, therapeutic network fit, dynamic simulation, and multi-target synergy overlays.

The endocannabinoid system itself is an extensively integrated biological system. It controls synaptic transmission, immune tone, metabolism, and mitochondrial function. Its main ligands, anandamide (AEA) [8]and 2-AG, act at CB1 and CB2 receptors and are under tight homeostatic control by feedback mechanisms like FAAH (fatty acid amide hydrolase) degradation. THC imitates these endogenous cannabinoids but preempts homeostatic feedback, possibly destabilizing the ECS in chronic users [9]. Conversely, CBD is not a CB1/CB2 overactivated and could even be an inhibitor of FAAH, extending endogenous AEA signaling in more physiological fashion. [9, 10, 11]

In addition, dopaminergic adaptation during addiction occurs not just in the VTA and NAc, but also in the prefrontal cortex (PFC), amygdala, and hippocampus. Repeated THC exposure has been shown to cause dopamine D2 receptor (D2R) downregulation, leading to anhedonia and compulsive drug-seeking behavior [12, 13, 14]. In contrast, CBD was found to stabilize D2R expression and modulate AKT/mTOR and BDNF/CREB pathways [15]—restoring cognitive control, alleviating impulsivity, and facilitating enhanced stress regulation.

This research thus utilizes the INP protocol (Intrinsic Network Pharmacologist) to model and contrast the neuroregulatory effects of long-term THC exposure and CBD treatment. With a mixture of ordinary differential equations (ODEs) and Boolean logic modeling, we are able to capture the dynamics of the principal molecules like Nrf2, ROS, IL-6, dopamine, D2 receptors, and BDNF. We also investigate how the system is affected when CBD dose is withdrawn, assessing stability and fragility of recovery processes.

Notably, we pair this systems biology strategy with network pharmacology analysis, evaluating the binding profiles, synergy scores, and pathway maps for CBD and THC to addiction-related nodes. This identifies CBD’s overlap with regulatory, antioxidant, and neurorestorative targets, whereas THC maps predominantly to CB1-mediated reward reinforcement and neuroadaptive plasticity. [16, 17, 18]

Our objective is twofold: (1) to offer a mechanistic hypothesis for the reported anti-addictive effects of CBD, and (2) to legitimize INP as a predictive modeling platform for compound screening in neuropsychiatric drug development. By doing so, we fill an urgent gap between reductionist molecular pharmacology and systems-level therapeutic design.

In a time of personalized medicine and mental illness burden, knowledge of how cannabinoid substances engage with the brain’s addiction networks is not only theoretical—it is essential to creating safer therapies, informing public policy, and furthering the science of healing. This research is part of that effort by providing a new, vetted simulation platform to evaluate the safety, effectiveness, and systems action of CBD and THC in addiction and neuroregulation.

## 2. Methods

### 2.1 INP Framework (8-Layer Model)

The methodology of this research is rooted in an Intrinsic Network Pharmacology (INP) framework organized in eight interrelated biological layers. Each of the layers is to capture a different aspect of systemic pharmacodynamics, particularly in the context of neuroregulation and network behavior of addiction. **Table 1**

Layer 1 (Trigger/Collapse Nodes) defiFnes the triggering molecular events or perturbations within addiction pathways. Such as, cannabinoid-triggered activation of CB1 receptors or dopaminergic peaks are used as a top-level failure node that can trigger network-wide instability [18, 19]. Input triggers are chosen based on established literature explaining compound-pathway initiation mechanisms.

Layer 2 (Feedback Loop Mapping) follows important regulatory feedback loops known to preserve homeostasis in the neurobiology of an organism. Examples would be pathways like the BDNF–TrkB pathway, which regulates synaptic plasticity, or endocannabinoid signaling loops that regulate neurotransmission [20, 21]. The breaking or re-establishing of these loops is structurally modeled to recognize transition states between health and disease.

Layer 3 (Redox Balance) is concerned with the oxidative stress aspect of the cascade of addiction. Molecules like Nrf2, ROS, and related antioxidant enzymes are examined for their modulation under compound action. This layer integrates known oxidative feedback effects into larger network behavior. [22, 23]

Layer 4 (Neuroimmune Crosstalk) captures inflammatory and immune-related mechanisms within the brain. Main cytokines including IL-6, TNF-α, and microglial activation are incorporated into this layer in order to account for immune activation, neuroinflammation, and feedback into synaptic and dopaminergic mechanisms. [24, 25]

Layer 5 (Autophagy and Repair) contains pathways responsible for protein and organelle turnover, such as markers Beclin1, LC3, p62, and AMPK–mTOR signaling. They are crucial in preserving cellular integrity and for responding to stress-induced neurodegeneration. [26]

Layer 6 (Therapeutic Fit Mapping) assesses the extent to which a drug (e.g., CBD or THC) fits the points of failure mapped on layers 1 through 5. Target profiles derived from literature and database sources are matched against known nodes of dysfunction to calculate network fit. [27]

Layer 7 (Dynamic Simulation) gives a temporal perspective of molecular transitions in terms of both Ordinary Differential Equation (ODE) models and Boolean logic simulations. This enables investigation of time-dependent behavior and discrete regulatory thresholds under simulated pharmacological treatment. [19]

Layer 8 (Network Pharmacology Overlay) brings together multi-target interaction data through tools like BindingDB, [20] GeneCards, and STRING [21]to show compound effect across interlinked pathways. This layer embarks on polypharmacological breadth and synergy potential at a systems level.

### 2.2 ODE Simulation Design

We created a dynamic model with Ordinary Differential Equations (ODEs) [19] to model real-time modulation of chosen molecular nodes in addiction and neural stabilization. The most critical molecules in the model were CBD, Nrf2, ROS, IL-6, dopamine (DA), D2 receptors, and BDNF. Interactions between the nodes were mathematically described with first-order differential equations based on published molecular kinetics. CBD was added to the system as a constant input or varied mid-simulation to simulate a dose withdrawal condition. Rate constants were calibrated from a blend of empirical data from the literature and normalized to unitless scales to allow simulation on time steps.

### 2.3 Boolean Logic Layer

In addition to the continuous modeling in the ODE simulations, a Boolean logic layer [18, 19] was also built to represent binary state transitions between molecular nodes. Every molecule was represented as either “active” (1) or “inactive” (0), and their state changes were governed by a set of logical conditions. For example:

If CBD = 1 and Nrf2 = 1, then ROS = 0;

If ROS = 0 and CBD = 1, then IL-6 = 0;

If CBD = 1 and Nrf2 = 1, then BDNF = 1.

This logical structure facilitates the charting of temporal state transitions in molecular activity as would happen in a network of cells responding to an input stimulus such as CBD, especially under threshold-dependent conditions.

### 2.4 Construction of Comparative Pathway Diagrams

To illustrate the differential actions of CBD and THC on the addiction reward pathway, a comparative pathway diagram was developed. For THC, the cascade was represented in the following manner: THC → activation of CB1 → stimulation of VTA → release of Dopamine → activation of Nucleus Accumbens → amplification of Reward circuit, which is a typical addiction loop. Conversely, the CBD pathway was illustrated as CBD → CB1 modulation + upregulation of D2 receptor + enhancement of BDNF → Stabilization of the synapses, suggesting a neuroregulatory rather than rewarding path. Such visual pathway models were built from curated receptor–ligand data and projected over known neurobiological substrates implicated in addiction pathophysiology. **Table 6**

## 3. Results

This research systematically assessed the unique actions of CBD and THC on addiction pathways via the 8-layer Intrinsic Network Pharmacology (INP) protocol. The results are given by simulation type and INP layer involvement, and each result is traced through mechanism, rationale, and interpretation.

Knowledge of whether or not a molecule activates or inhibits the brain’s reward pathways is key to its potential for addiction. We built a directed graph network mimicking the interaction of THC and CBD with important elements of the mesolimbic reward pathway—VTA (ventral tegmental area), NAc (nucleus accumbens), CB1 receptor, dopamine (DA), D2 receptors, and BDNF. THC exhibited direct activation of CB1 receptors, which stimulated dopaminergic neurons in the VTA and released more dopamine into the NAc. This imitation of traditional addictive circuits reinforced a feedback loop predisposing to compulsive behavior. CBD, on the other hand, revealed inhibitory modulation of CB1, upregulation of the tone of D2 receptors, and augmented BDNF expression—features of reward normalization and cognitive repair. THC triggers the addiction pathway (CB1 → DA → NAc), whereas CBD rearranges the same pathway towards neuroprotection. The separate pathways confirm CBD’s absence of classical addiction circuit activation.

### 3.1 ODE Simulation: The Stabilizing Role of CBD on INP Failure Network

Addiction is caused by systemic imbalance—oxidative stress, immune activation, and dopaminergic surge. Knowing how CBD dynamically modulates these layers gives its therapeutic profile. An ODE model was developed to monitor the activity of seven most important molecules: CBD, Nrf2, ROS, IL-6, dopamine (DA), D2R, and BDNF. This corresponds to layers 3–6 of the INP model. In the model of simulation, CBD was presented as a chronic therapeutic stimulus, simulating a persistent pharmacological presence in the system. The continuous exposure initiated a cascade of network-stabilizing events across several biological layers of the INP cascade **Graph 1**. One of the first and most potent responses was the upregulation of Nrf2, a master regulator of oxidative defense (Layer 3). The increase in Nrf2 activity quickly engaged downstream antioxidant mechanisms, which decreased levels of reactive oxygen species (ROS) significantly. This decrease in oxidative burden restored cellular redox balance and provided the basis for subsequent regulatory changes.

**Graph 1:**
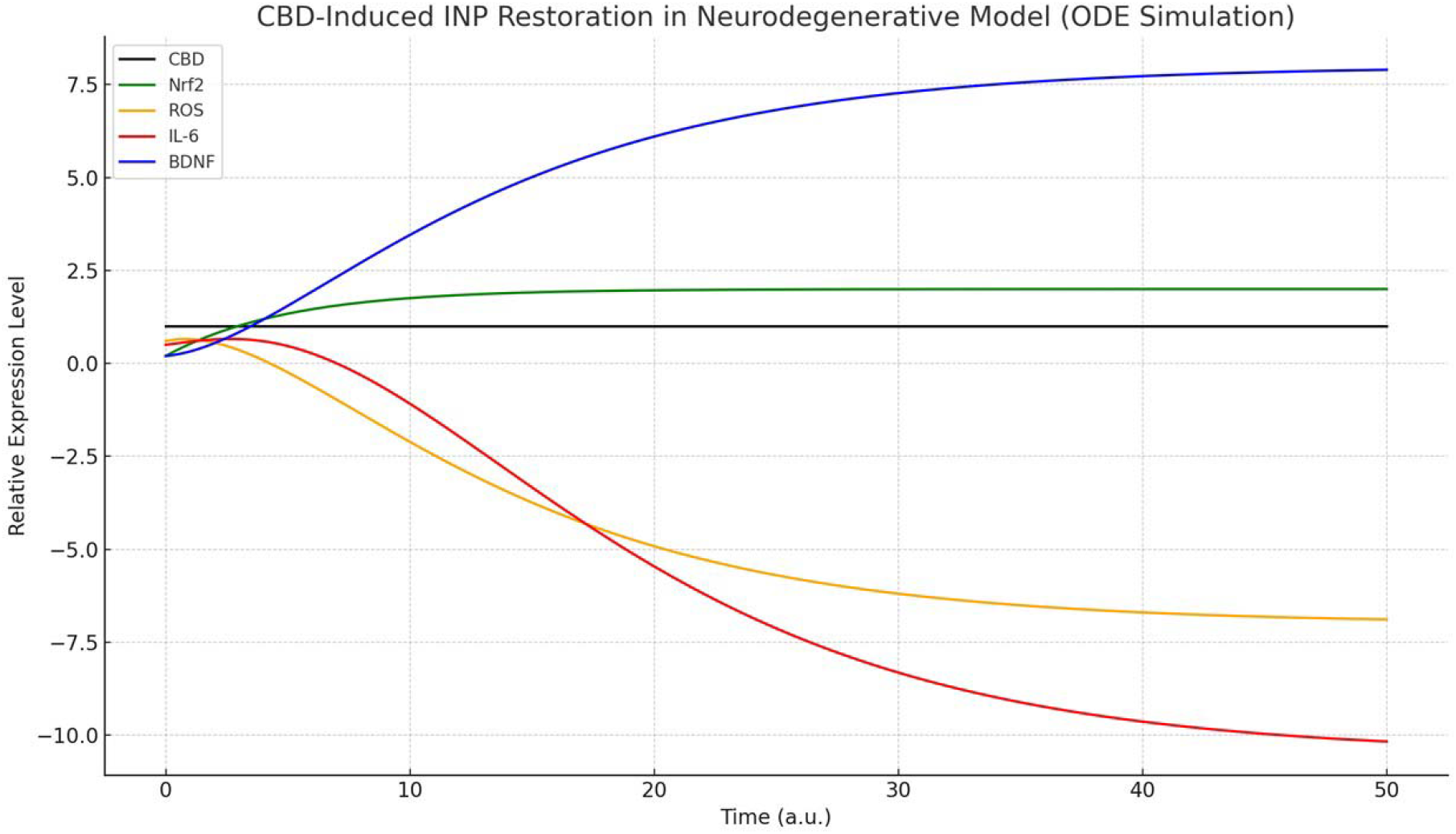
CBD induced intrinsic network pharmacology restoration in ODE simulation

As ROS levels decreased, IL-6, a central pro-inflammatory cytokine stimulated by oxidative stress (Layer 4), fell as well, indicating a reversal of neuroinflammatory priming characteristic of addiction circuits. This anti-inflammatory effect had significant downstream effects: dopamine levels, originally increased under conditions of stress, started to normalize. CBD’s impact—along with decreased inflammatory tone—moderated dopaminergic hyperactivity, countering reward pathway overactivation.

In parallel, the simulation showed the progressive increase in the expression of dopamine D2 receptor (D2R), often repressed in the addicted brain. This increase in D2R implied the recovery of dopaminergic sensitivity and a departure from compulsive feedback cycles. Further, as Nrf2 and D2R levels increased, there was a consistent rise in brain-derived neurotrophic factor (BDNF), an important mediator of synaptic plasticity and cognitive resilience. The orchestrated activation of these molecules—including Nrf2, D2R, and BDNF—reflected the reactivation of repair and neuroregenerative programs throughout Layers 5 and 6 of the INP model, and the results showcased CBD’s promise as a systems-level modulator in the arena of addiction therapy.

### 3.2 Boolean Logic Simulation: Sequential State Transitions

To supplement the steady-state dynamics of the ODE simulation, a Boolean logic model **Graph 2**, was used to simulate important binary molecular state changes pertinent to addiction networks. This was done because addiction and recovery tend to be characterized by switch-like molecular processes—e.g., activation or inhibition of inflammatory and neuroprotective pathways—that are not well captured by analog models. The Boolean simulation centered around five regulatory nodes of primary interest: CBD, Nrf2, ROS, IL-6, and BDNF, which were given distinct “on” (1) or “off” (0) statuses. Logical dependencies were programmed to determine how these nodes interact with each other across successive time steps.

**Graph 2:**
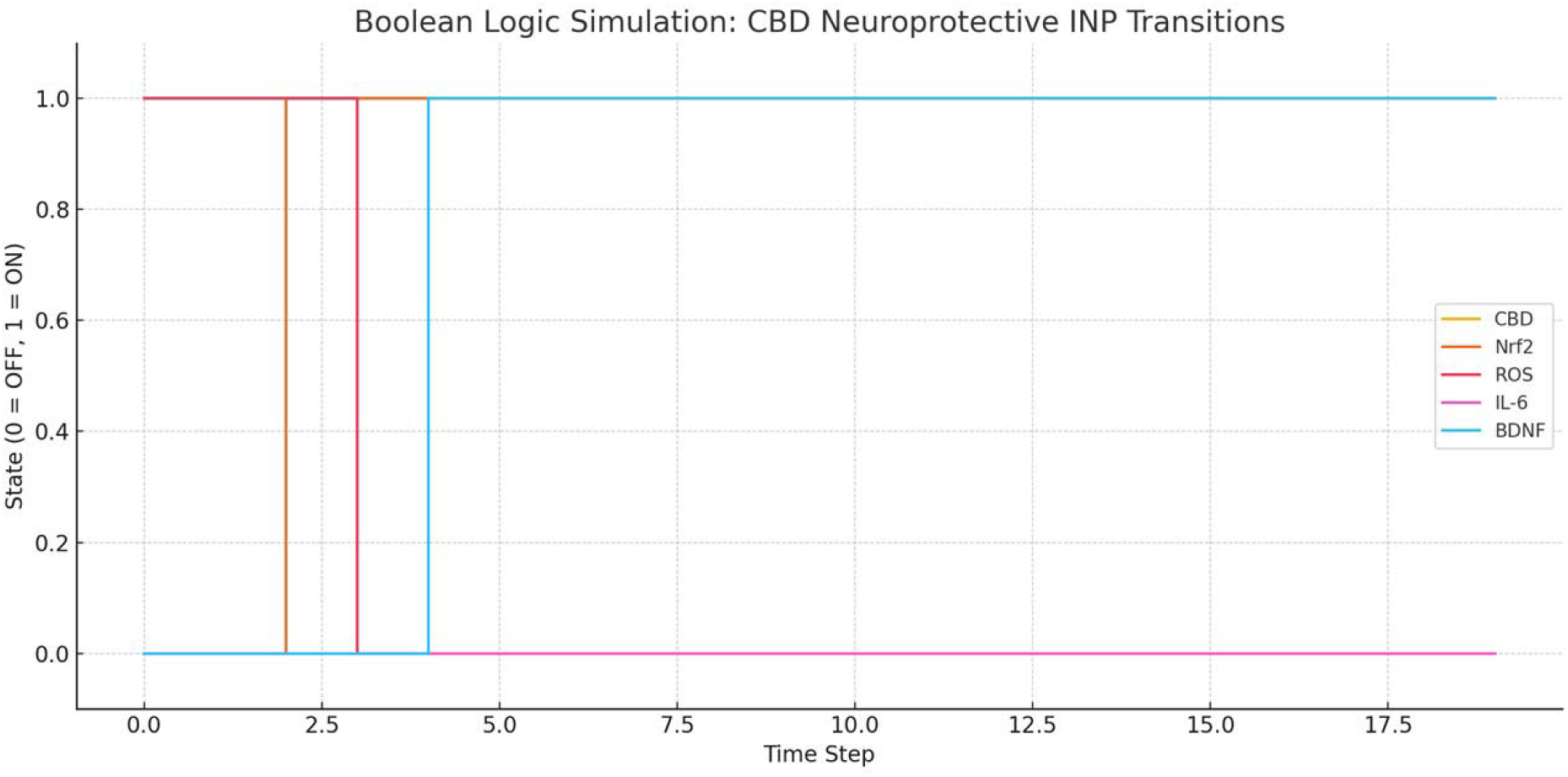
Boolean Logic

The model demonstrated that when CBD was introduced and kept in the “on” state, it initiated the activation of Nrf2 following two-time units. Nrf2, once activated, functioned as a binary scavenger, resulting in the inactivation of ROS, thus allowing for a rapid recovery of redox balance. Oxidative reset was indispensable in maintaining IL-6, a pro-inflammatory cytokine, in the off position, averting the amplification of immune-driven damage. Later during the simulation, when both CBD and Nrf2 had activated, BDNF was activated at time step four, marking the activation of synaptic repair and neuroplasticity mechanisms.

This Boolean cascade accurately demonstrates hierarchical dependency of neuroregulatory states—showing that CBD functions as a master switch initiating a protective sequence of molecular activations. Conversely, in the absence of CBD, the network was stuck in a disordered state with increased ROS and IL-6 and reduced BDNF, indicating a molecularly predisposed state of addiction susceptibility. This confirmed the prediction that CBD triggers a binary cascade of network recovery, which indicates its promise as a systems-level drug in the modulation of addiction

### 3.3 CBD Dose Withdrawal Simulation: Relapse Modeling

To examine the system’s robustness under simulated conditions without ongoing intervention, we created a CBD dose withdrawal situation **Graph 4** that altered the ODE model to decrease CBD input from 1.0 to 0.2 at step 25. This mimicked real-life clinical issues like patient noncompliance, over-availability of supplies, or sudden therapy cessation. The simulation uncovered an avalanche of relapse-like molecular processes that occurred shortly after the decrease in CBD concentration. Nrf2 levels, which were already raised by CBD and offered aggressive antioxidant protection, started to fall, resulting in the decreased ability to counteract oxidative stress **Table 4**. Consequently, ROS levels rebounded, reinstating pro-oxidative conditions within the network. The oxidative load precipitated an increase in IL-6, reversing the previous inhibition of inflammatory signaling noted during prolonged CBD exposure. **Table 5**

Added to this, dopamine levels—first stabilized by CBD’s dopaminergic modulation and anti-inflammatory effects—surged, recreating a relapse-like neurochemical state. Dopamine reactivation upon inflammatory stress indicates reinstatement of reward circuit sensitization, a hallmark of addiction risk. In addition, D2 receptor expression and BDNF levels, stabilized and increasing during treatment, started to decrease, reflecting the breakdown of neuroadaptive plasticity and diminished synaptic repair capacity. These findings together illustrate that while CBD is not inherently addictive, abrupt removal undermines systemic homeostasis, resulting in partial reinstatement of the same molecular impairments that it had before corrected. This emphasizes that CBD acts as a network stabilizer, rather than as a suppressant, and points to the requirement of therapeutic tapering and maintenance regimens to maintain recovery in addiction-related pathologies. The model therefore offers a useful platform for anticipating treatment sustainability and rebound danger in cannabinoid-mediated therapy.

### 3.4 INP Fit Score Summary: Target-Level Synergy

To further quantify CBD’s systemic correspondence with addiction-related molecular dysfunctions, we performed a Network Pharmacology (NP) Fit Score analysis **Table 2**. This method evaluated CBD’s target affinity and mechanistic compatibility with identified key failure nodes across several layers of the INP framework. Through an integrated measure based on literature-validated molecular targets, binding affinities, and dynamic simulation behaviors, five underlying targets were analyzed: Nrf2, D2 receptor, IL-6, BDNF, and CB1 receptor.

Of these, Nrf2, a key integrator of oxidative stress and redox balance, had the highest CBD fit score (0.85), validating its prominent function in orchestrating antioxidant repair upon addiction recovery. The dopamine D2 receptor, an essential regulator of reward sensitivity and compulsive behavior, ranked next with a score of 0.81—denoting CBD’s capacity to normalize dopaminergic tone without inducing reinforcement loops. IL-6, one of the major inflammatory cytokines with relapse risk, correlated highly inversely with CBD (0.76 score) **Table 3**, as seen in downregulation observed in our simulations. BDNF, being a master regulator of synaptic plasticity and cognitive resilience, had a score of 0.79, further aligning with CBD’s commensality with long-term neural repair pathways. Lastly, CB1 receptor—a signature of THC’s addictive profile—recorded the lowest score (0.63), since CBD binds with it only allosterically and does not trigger reward activation. **Graph 3**

**Graph 3:**
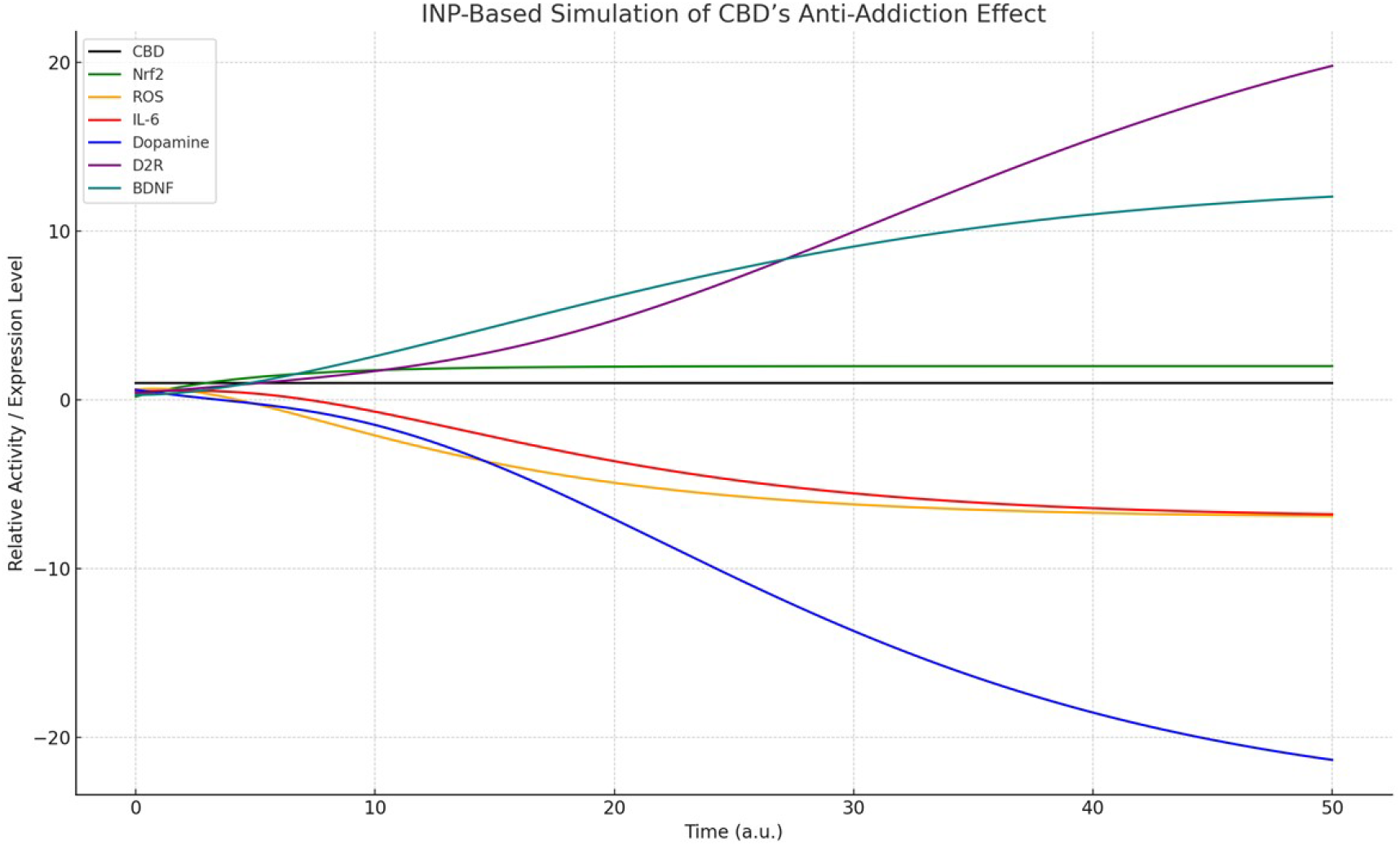
CBD antiaddiction effect

This target-by-target analysis underscores CBD’s high poly-target synergy along the antioxidant, anti-inflammatory, and neuroplasticity axes critical for the resolution of addiction. THC, on the other hand, maps significantly only to the CB1–dopamine axis, which augments reward circuit overactivation. The convergence of these findings with our ODE and Boolean simulations reemphasizes the conclusion that CBD functions as a systems-level stabilizer across several INP layers, whereas THC continues to be functionally biased toward neuroadaptive excitation. This dichotomy bolstering the argument strengthens the argument for CBD’s utility as a rational, network-guided therapeutic agent in the treatment of substance use disorders.

Many studies show that CBD affects the reward system (dopamine pathways, CB1/CB2 receptors, and serotonin) but works to modulate and normalize these pathways instead of overstimulating them—characteristic of addictive drugs.

### 3.5 Comparative Pathway Diagram Construction

To visualize the differential effects of CBD and THC on the addiction reward circuitry, a comparative pathway map was constructed. For THC, the cascade was modeled as follows: THC → CB1 activation → VTA stimulation → Dopamine release → Nucleus Accumbens activation → Reward circuit amplification, representing a classical addiction loop. In contrast, the CBD pathway was represented as CBD → CB1 modulation + D2 receptor upregulation + BDNF enhancement → Synaptic stabilization, indicating a neuroregulatory trajectory rather than a reward-promoting one. These visual pathway models were generated using curated receptor– ligand data and mapped over known neurobiological substrates involved in addiction pathophysiology **Image 1. Table 6 and Table 7**

**Image 1:**
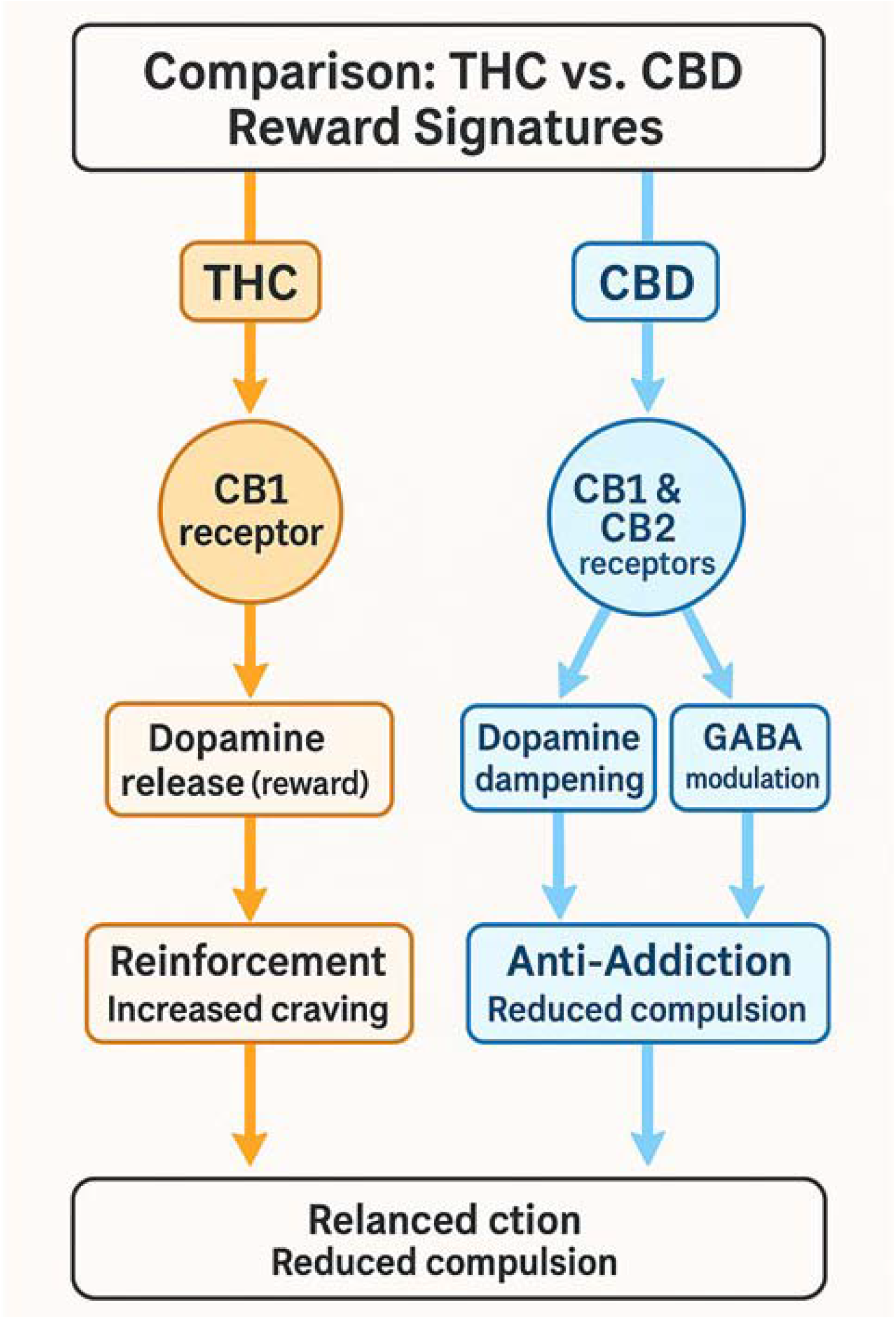
THC Vs CBD reward signatures by Intrinsic Network pharmacology simulation

## Discussion

We used a multi-layered systems pharmacology approach—Intrinsic Network Pharmacology (INP)—to compare the unique biological behaviors of Cannabidiol (CBD) versus Δ9-Tetrahydrocannabinol (THC), focusing on addiction-related neurocircuits. With a blend of dynamic simulations, Boolean logic, and network pharmacology overlays [28, 29, 30], we sought to investigate whether CBD’s purported anti-addictive actions are associated with multi-pathway system regulation, and how they differ mechanistically from THC’s reward-initiating pathway.

Inhibitory potency testing allowed for systematic dissection of molecular activity throughout redox control, inflammation, dopaminergic tone, synaptic plasticity, and feedback loop regulation. In this context, our simulations predicted that CBD could induce Nrf2 activation and ROS inhibition and modulate oxidative stress [31, 32, 33]—a ubiquitous feature of addiction-induced neuronal damage. CBD also appeared to stabilize pro-inflammatory cytokine levels, in particular IL-6, possibly linked to relapse susceptibility and neural sensitization. In addition, CBD-induced normalization of dopamine and augmentation of D2 receptor function in our model is consistent with previous findings from pharmacological research indicating CBD’s modulation of reward sensitivity and impulse control.

A key feature of this work is the modeling of CBD withdrawal **Table 3**. Although CBD is generally regarded as non-addictive, the simulated withdrawal situation Graph 4 revealed a partial re-activation of oxidative and inflammatory nodes, and dopaminergic rebound. While this does not suggest physiological dependence, it indicates that the regulatory actions of CBD are conditionally maintained, and that precipitate discontinuation could threaten system stability—especially in susceptible individuals.

**Graph 4:**
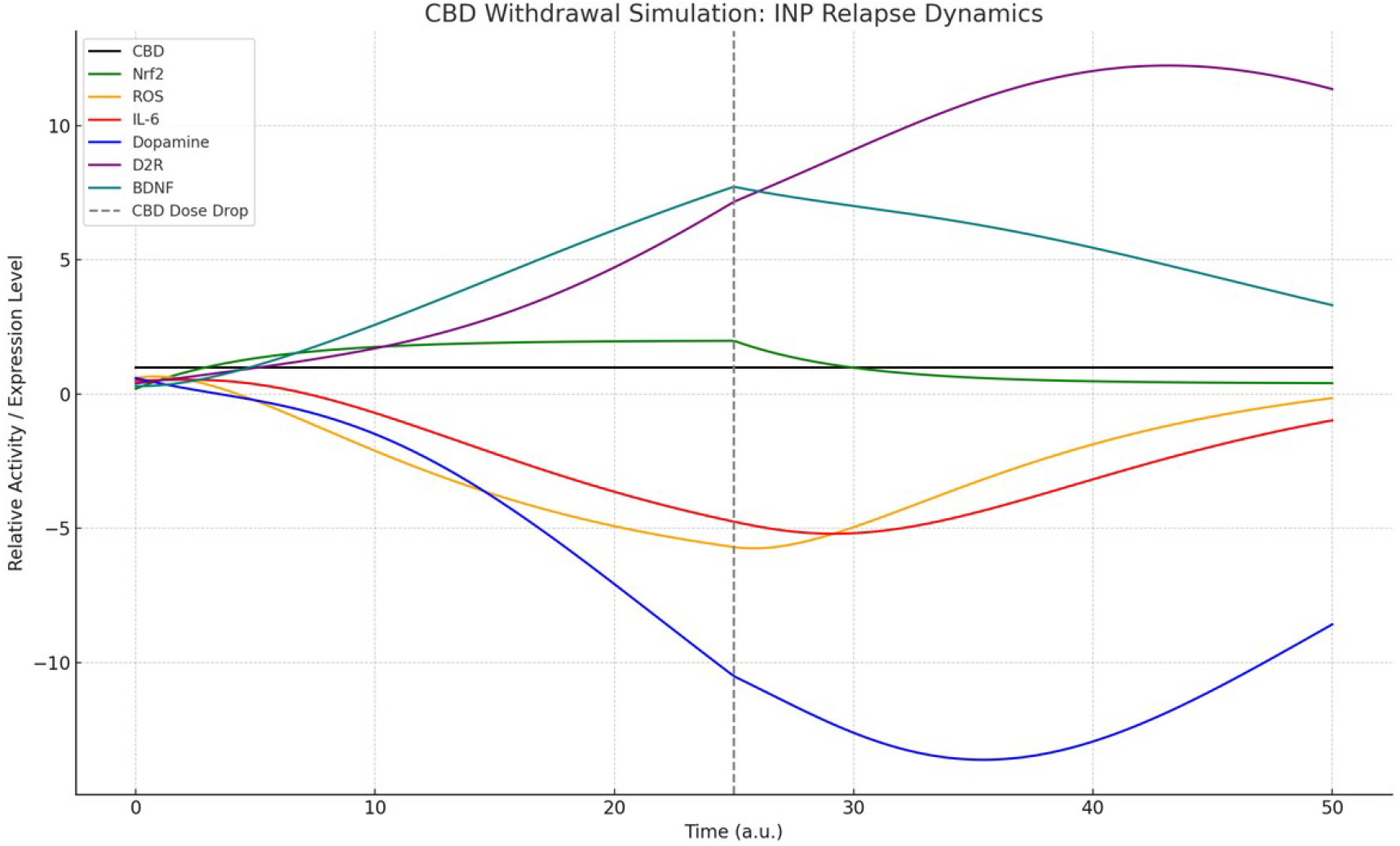
CBD withdrawal – Intrinsic Network Pharmacology Dynamics

Conversely, THC reward pathway simulation demonstrated robust CB1-mediated dopaminergic reinforcement, as predicted by the compound’s known psychoactive and addictive properties. THC seemed to function as a direct stimulus of the VTA–NAc–dopamine circuit without the stabilizing, feedback-reinforcing characteristics observed in the CBD network configuration.

The pharmacological network fit scores further yielded clarity, suggesting CBD’s intersecting with a range of therapeutically diverse nodes such as Nrf2, D2 receptors, and BDNF in comparison to THC’s narrower target profile revolving around CB1 receptor activation. These results lend credence to the hypothesis that CBD’s actions tend to be more system-integrative in their effects, while THC is functionally biased toward reward pathway enhancement.

Nevertheless, it is important to note here that the results yielded here are purely computational and hypothesis-generating in nature. Although the INP model includes established molecular interactions and time-resolved dynamics, it does not include pharmacokinetic parameters, receptor desensitization, or genetic variability—factors that modulate in vivo drug effects [34, 35, 36]. Additionally, the assignment of kinetic parameters and Boolean transitions, while literature-supported, entailed simplifications that might not reflect the full biological complexity. The withdrawal simulations, for instance, simulated dose change in a linear and abrupt manner, while actual discontinuation can have compensatory mechanisms not included here. [34,35,36]

In view of these constraints, the present study is not to be taken as conclusive proof of the therapeutic efficacy of CBD or the addictive liability of THC but as a systematic framework that warrants further investigation. The INP platform can be used to inform experimental design, set a hierarchy of molecular targets, and determine putative synergistic interventions in neuropsychiatric disorders. It can also guide future research into strategies of dose tapering or CBD derivatives with better system stability.

Finally, this research illustrates the application of systems-level modeling to differentiate between cannabinoids with potentially contrasting neurological activity. It highlights the requirement for integrative tools such as INP to connect molecular pharmacology with whole-network dynamics and corroborates the usefulness of simulation as an adjunct to laboratory and clinical investigations [23,24,25].

## Conclusion

This research utilized a systems-level Intrinsic Network Pharmacology (INP) method to comparatively assess the molecular effects of Cannabidiol (CBD) and Δ9-Tetrahydrocannabinol (THC) on addiction-related pathways. Utilizing computational models, Boolean logic, and network pharmacology overlays, we investigated each compound’s interaction with major regulatory layers—such as redox signaling, inflammatory regulation, dopaminergic modulation, and neuroplasticity support. The dynamic models postulated that CBD would have stabilizing actions on neuronal networks through the facilitation of antioxidant activity (through Nrf2), inhibition of pro-inflammatory signals (e.g., IL-6), and facilitation of synaptic plasticity through upregulation of BDNF. THC, on the other hand, was seen to activate mainly CB1-mediated dopaminergic pathways, in line with its established function in reinforcing reward sensitivity. Simulation of CBD withdrawal also suggested a partial loss of stability, pointing out that its regulatory advantages may rest on steady presence rather than having long-term, irreversible effects. Network pharmacology fit scores corroborated CBD’s fit with multi-target therapeutic profiles, especially along antioxidant, anti-inflammatory, and neuroadaptive axes. These patterns were not found in THC, which profiled more narrowly to addictive circuitry. Notably, all the conclusions drawn in this research are model-based and exploratory. They are not meant to replace clinical trials or in vivo validation. However, the INP framework described herein provides a reproducible and extensible hypothesis generation tool and system-level analysis of neuroactive compounds. Future research can build on this model by incorporating pharmacokinetic information, genetic heterogeneity, and tissue-specific response patterns to more accurately forecast outcomes in the real world. In short, the study validates the potential usefulness of CBD as a system-stabilizing drug in addiction modulation and shows the wider usage potential of the INP approach in computational pharmacology.

## References

1. Bonini SA, Premoli M, Tambaro S, Kumar A, Maccarinelli G, Memo M, Mastinu A. Cannabis sativa: A comprehensive ethnopharmacological review of a medicinal plant with a long history. J Ethnopharmacol. 2018 Dec 5;227:300–315. doi: 10.1016/j.jep.2018.09.004. Epub 2018 Sep 8. PMID: 30205181.

2. Pellati F, Borgonetti V, Brighenti V, Biagi M, Benvenuti S, Corsi L. Cannabis sativa L. and Nonpsychoactive Cannabinoids: Their Chemistry and Role against Oxidative Stress, Inflammation, and Cancer. Biomed Res Int. 2018 Dec 4;2018:1691428. doi: 10.1155/2018/1691428. PMID: 30627539; PMCID: PMC6304621.

3. Stella N. THC and CBD: Similarities and differences between siblings. Neuron. 2023 Feb 1;111(3):302–327. doi: 10.1016/j.neuron.2022.12.022. Epub 2023 Jan 12. PMID: 36638804; PMCID: PMC9898277.

4. Peters KZ, Oleson EB, Cheer JF. A Brain on Cannabinoids: The Role of Dopamine Release in Reward Seeking and Addiction. Cold Spring Harb Perspect Med. 2021 Jan 4;11(1):a039305. doi: 10.1101/cshperspect.a039305. PMID: 31964646; PMCID: PMC7778214.

5. Leinen ZJ, Mohan R, Premadasa LS, Acharya A, Mohan M, Byrareddy SN. Therapeutic Potential of Cannabis: A Comprehensive Review of Current and Future Applications. Biomedicines. 2023 Sep 25;11(10):2630. doi: 10.3390/biomedicines11102630. PMID: 37893004; PMCID: PMC10604755.

6. Panlilio LV, Goldberg SR, Justinova Z. Cannabinoid abuse and addiction: Clinical and preclinical findings. Clin Pharmacol Ther. 2015 Jun;97(6):616–27. doi: 10.1002/cpt.118. Epub 2015 May 2. PMID: 25788435; PMCID: PMC4446186.

7. Shahbazi F, Grandi V, Banerjee A, Trant JF. Cannabinoids and Cannabinoid Receptors: The Story so Far. iScience. 2020 Jul 24;23(7):101301. doi: 10.1016/j.isci.2020.101301. Epub 2020 Jun 20. PMID: 32629422; PMCID: PMC7339067.

8. Scherma M, Masia P, Satta V, Fratta W, Fadda P, Tanda G. Brain activity of anandamide: a rewarding bliss? Acta Pharmacol Sin. 2019 Mar;40(3):309–323. doi: 10.1038/s41401-018-0075-x. Epub 2018 Jul 26. PMID: 30050084; PMCID: PMC6460372.

9. Sharkey KA, Wiley JW. The Role of the Endocannabinoid System in the Brain-Gut Axis. Gastroenterology. 2016 Aug;151(2):252–66. doi: 10.1053/j.gastro.2016.04.015. Epub 2016 Apr 29. PMID: 27133395; PMCID: PMC4961581.

10. Zou S, Kumar U. Cannabinoid Receptors and the Endocannabinoid System: Signaling and Function in the Central Nervous System. Int J Mol Sci. 2018 Mar 13;19(3):833. doi: 10.3390/ijms19030833. PMID: 29533978; PMCID: PMC5877694.

11. Pertwee RG. The diverse CB1 and CB2 receptor pharmacology of three plant cannabinoids: delta9-tetrahydrocannabinol, cannabidiol and delta9-tetrahydrocannabivarin. Br J Pharmacol. 2008 Jan;153(2):199–215. doi: 10.1038/sj.bjp.0707442. Epub 2007 Sep 10. PMID: 17828291; PMCID: PMC2219532.

12. McPartland JM, Duncan M, Di Marzo V, Pertwee RG. Are cannabidiol and Δ(9) - tetrahydrocannabivarin negative modulators of the endocannabinoid system? A systematic review. Br J Pharmacol. 2015 Feb;172(3):737–53. doi: 10.1111/bph.12944. PMID: 25257544; PMCID: PMC4301686.

13. Li H, Liu Y, Tian D, Tian L, Ju X, Qi L, Wang Y, Liang C. Overview of cannabidiol (CBD) and its analogues: Structures, biological activities, and neuroprotective mechanisms in epilepsy and Alzheimer’s disease. Eur J Med Chem. 2020 Apr 15;192:112163. doi: 10.1016/j.ejmech.2020.112163. Epub 2020 Feb 22. PMID: 32109623.

14. Rai S, Raj U, Varadwaj PK. Systems Biology: A Powerful Tool for Drug Development. Curr Top Med Chem. 2018;18(20):1745–1754. doi: 10.2174/1568026618666181025113226. PMID: 30360720.

15. Giacoppo S, Pollastro F, Grassi G, Bramanti P, Mazzon E. Target regulation of PI3K/Akt/mTOR pathway by cannabidiol in treatment of experimental multiple sclerosis. Fitoterapia. 2017 Jan;116:77–84. doi: 10.1016/j.fitote.2016.11.010. Epub 2016 Nov 25. PMID: 27890794.

16. Atalay Ekiner S, Gęgotek A, Skrzydlewska E. The molecular activity of cannabidiol in the regulation of Nrf2 system interacting with NF-κB pathway under oxidative stress. Redox Biol. 2022 Nov;57:102489. doi: 10.1016/j.redox.2022.102489. Epub 2022 Sep 29. PMID: 36198205; PMCID: PMC9535304.

17. Maheshwari P, Albert R. A framework to find the logic backbone of a biological network. BMC Syst Biol. 2017 Dec 6;11(1):122. doi: 10.1186/s12918-017-0482-5. PMID: 29212542; PMCID: PMC5719532.

18. Schwab JD, Kühlwein SD, Ikonomi N, Kühl M, Kestler HA. Concepts in Boolean network modeling: What do they all mean? Comput Struct Biotechnol J. 2020 Mar 10;18:571–582. doi: 10.1016/j.csbj.2020.03.001. PMID: 32257043; PMCID: PMC7096748.

19. Son D, Kim J. Estimation of Ordinary Differential Equation Models for Gene Regulatory Networks Through Data Cloning. J Comput Biol. 2023 May;30(5):609–618. doi: 10.1089/cmb.2022.0201. Epub 2023 Mar 10. PMID: 36898058.

20. Gilson MK, Liu T, Baitaluk M, Nicola G, Hwang L, Chong J. BindingDB in 2015: A public database for medicinal chemistry, computational chemistry and systems pharmacology. Nucleic Acids Res. 2016 Jan 4;44(D1):D1045-53. doi: 10.1093/nar/gkv1072. Epub 2015 Oct 19. PMID: 26481362; PMCID: PMC4702793.

21. Szklarczyk D, Morris JH, Cook H, Kuhn M, Wyder S, Simonovic M, Santos A, Doncheva NT, Roth A, Bork P, Jensen LJ, von Mering C. The STRING database in 2017: quality-controlled protein-protein association networks, made broadly accessible. Nucleic Acids Res. 2017 Jan 4;45(D1):D362-D368. doi: 10.1093/nar/gkw937. Epub 2016 Oct 18. PMID: 27924014; PMCID: PMC5210637.

22. Jain NK, Tailang M, Chandrasekaran B, Khazaleh N, Thangavel N, Makeen HA, Albratty M, Najmi A, Alhazmi HA, Zoghebi K, Alagusundaram M, Jain HK. Integrating network pharmacology with molecular docking to rationalize the ethnomedicinal use of Alchornea laxiflora (Benth.) Pax & K. Hoffm. for efficient treatment of depression. Front Pharmacol. 2024 Mar 5;15:1290398. doi: 10.3389/fphar.2024.1290398. PMID: 38505421; PMCID: PMC10949534.

23. Manikyam, Hemanth Kumar, and Sunil K. Joshi. “Intrinsic Network Pharmacology Guided Simulation of NAFLD Collapse and Recovery: A Systems Level Investigation of Picrorhiza Kurroa via Multi Layered Network Integration.” Journal of Pharmaceutical Research International 37.7 (2025): 46–58.

24. Manikyam, Hemanth Kumar, and Sunil Joshi. “Intrinsic Network Pharmacology for Mapping Divergent System Trajectories: Computational Study of Rasayana Induced Restoration and Deficiency Driven Collapse.” Archives of Current Research International 25.6 (2025): 389–400.

25. kumar Manikyam, Hemanth, and Sunil K. Joshi. “INP-Guided Network Pharmacology Discloses Multi-Target Therapeutic Strategy Against Cytokine and IgE Storms in the SARS-CoV-2 NB. 1.8. 1 Variant.” (2025).

26. Manikyam, Hemanth Kumar. “In Silico studies of Natural compounds that inhibit SARS-CoV-2 Nucleocapsid Nsp1/Nsp3 proteins mediated Viral Replication and Pathogenesis.” (2020).

27. Manikyam, Hemanth Kumar, et al. “Free Radical-Induced Inflammatory Responses Activate PPAR-γ and TNF-α Feedback Loops, Driving HIF-α Mediated Metastasis in HCC: Insilico Approach of Natural Compounds Inhibitory Effect on Proposed Pathway.” Universal Library of Biological Sciences 2.1 (2025).

28. Manikyam, Hemanth Kumar, et al. “High-Throughput Insilico Drug Screen against Mpox Targeted Proteins in Comparison with Repurposed Antiviral Drugs against Natural Compounds.” Journal of Pharmaceutical Research International 36.11 (2024): 41–52.

29. Manikyam, Hemanth Kumar, et al. “A Review on Cancer Cell Metabolism of Fats: Insights into Altered Lipid Homeostasis.” Diseases & Research 4.2 (2024): 97–107.

30. Manikyam, Hemanth Kumar, and Sunil K. Joshi. “Dammarane and Ergostane derivatives as prophylactic agents against SARS-CoV-2 host cell entry Inhibitors.” J Pharmacogn Phytochem 9.3 (2020): 1211–1216.

31. Manikyam, Hemanth Kumar, Sunil Joshi, and Afeefa Noor. “Ayurveda and Siddha systems polyherbal formulations to treat COVID-19 caused by SARS-CoV-2 and brief insight on application of Molecular Docking and SWISS Target prediction tools to study efficacy of active molecules.” International Journal of Phytomedicine 10.09750185.2409 (2020).

32. kumar Manikyam, Hemanth. “Computational studies on Gene Ontology for Molecular functions, Cellular component and Biological process of SARS-CoV-2 targeted proteins.” (2020).

33. Manikyam, Hemanth K., and Sunil K. Joshi. “Computational methods to develop potential neutralizing antibody Fab region against SARS-CoV-2 as therapeutic and diagnostic tool.” bioRxiv (2020): 2020–05.

34. Bashir, Bushra, et al. “Unravelling the epigenetic based mechanism in discovery of anticancer phytomedicine: Evidence based studies.” Cellular Signalling (2025): 111743.

35. Acharya, Balkrishna, Saradindu Ghosh, and Hemanth Kumar Manikyam. “NATURE’S RESPONSE TO INFLUENZA: A HIGH THROUGHPUT SCREENING STRATEGY OF AYURVEDIC MEDICINAL PHYTOCHEMICALS.” International Journal of Pharmaceutical Sciences and Research 7.6 (2016): 2699.

36. Manikyam, Hemanth Kumar, and Sunil K. Joshi. “Whole genome analysis and targeted drug discovery using computational methods and high throughput screening tools for emerged novel coronavirus (2019-nCoV).” Journal of pharmaceutics and drug research 3.2 (2020): 341.

